# Roads as conduits of taxonomic, functional and phylogenetic degradation in caatinga vegetation

**DOI:** 10.1101/2020.03.27.012286

**Authors:** Nayara Mesquita Mota, Markus Gastauer, Juan Fernando Carrión, João Augusto Alves Meira-Neto

## Abstract

Road networks cause disturbances that can alter the biodiversity and the functioning of the Caatinga ecosystems. We tested the hypotheses that (i) Caatinga vegetation near roads is less taxonomically, functionally and phylogenetically diverse, (ii) phylogenetically and functionally more clustered than vegetation further from roads, (iii) plant traits associated with herbivory deterrence are conserved within the phylogenetic lineages, and (iv) Caatinga vegetation near roads selects for disturbance-related traits. We sampled herbaceous and woody component of vegetation in four plots near roads and four plots further from roads to test these hypothesis. Sampled species were classified according to their resprouting capacity, nitrogen fixation, succulence/spines, urticancy/toxicity, lifeform, endozoochory, maximum height and maximum diameter, before we calculated the taxonomic, functional and phylogenetic diversity of plant communities. Species richness, taxonomic, functional and phylogenetic diversities were lower in plots close to the roads, confirming roads as sources of disturbances. The phylogenetic structure of the Caatinga vegetation near roads was clustered, indicating environmental filtering by herbivory as the main pervasive disturbance in Caatinga ecosystems, since traits related to herbivory deterrence were conserved within phylogenetic lineages and were filtered in near roads. Thus, roads should be considered degradation conduits causing taxonomic, phylogenetic and functional impoverishment of Caatinga vegetation.

## Introduction

Road networks change the functioning of the ecosystems by modifying soil properties, hydrological regimes, ecological flows, and, especially, amplifying disturbances in different ways. (Forman and Alexander, 1998; Laurance et al., 2009). If we understand disturbance as biomass loss (Tilman, 1988), disturbances reduce partially or totally the plant biomass of an ecosystem altering its original biodiversity and functioning (Hooper et al., 2005). Road networks serve as conduits for disturbances and functions (Christen and Matlack, 2009; Forman and Alexander, 1998; Forman and Deblinger, 2000) that can be guided and amplified on roadsides. In general, such disturbances are associated with clearcutting, extraction of firewood, soil movement, or grazing of domestic animals (Forman, 1995; Spooner et al., 2004; Watkins et al., 2003). Thus, roads and associated disturbances can alter habitats for the benefit of some plant species, including invasive species, as well as can alter habitats to the detriment of native plant species (Heringer et al., 2019b, 2019a; Lehmann et al., 2017; Spooner et al., 2004). As a result, disturbances associated with roads result in changes of ecosystem functioning due to increased habitat loss, fragmentation, and creation of novel habitats (Karim and Mallik, 2008; Rentch et al., 2005; Santos and Tabarelli, 2002; Trombulak and Frissell, 2000). One of the most altered Neotropical biomes by human disturbances is the Caatinga, a Brazilian Northeastern semiarid vegetation also referred as dry forest (Dryflor et al., 2016; Santos and Tabarelli, 2002). As disturbances are conducted, and amplified by roads, Caatinga near and far from roads should differ in terms of diversity, structure, and functioning.

The consequences of disturbances on vegetation depends on the regime and on the affected ecosystems. Partial cut as well as clear-cut disturbances in dry forests drive to loss of aboveground biomass that remains for more than a decade (Niño et al., 2014). Chronic disturbances as selective cutting and extraction of firewood cause phylogenetic impoverishment throughout age structure of plants in Caatinga (Ribeiro et al., 2016).

The most pervasive chronic disturbance of Caatinga is the overgrazing of domestic animals, especially goats (Leal et al., 2005). Herbivory by goats, functions as a selective pressure that may affect the abundance and distribution of the Caatinga flora, since it can reduce the richness of succulent fruit species, the richness of geophytes and the richness of nitrogen fixing species (Moolman and Cowling, 1994; Severson and DeBano, 1991). Overgrazing by domestic animals causes changes in ecosystems because of selective herbivory on seedlings causing expansion of non-palatable species (Bucher 1987), and can cause various types of changes to the functioning of the Caatinga, such as impairing the regeneration of arboreal species, preventing the dispersion of fruits and seeds, decreasing seedling survival, and limiting ecosystem productivity (Leal et al., 2003).

The consequences of disturbances on vegetation depends on the regime and on the affected ecosystems. Partial cut as well as clear-cut disturbances in dry forests drive to loss of aboveground biomass that remains for more than a decade. Chronic disturbances as selective cutting and extraction of firewood cause phylogenetic impoverishment throughout age structure of plants in Caatinga (Ribeiro et al., 2016).

Functional, and phylogenetic diversities, and structures are useful tools for predicting the ecological consequences of disturbances, and other anthropogenic changes (Cadotte et al., 2009; Edwards et al., 2007; Petchey and Gaston, 2006), including disturbance by herbivory. For instance, decreased functional diversity might indicate that some of the resources in an ecosystem are no longer fully available (Mason et al., 2005). The ecosystem resources would not be available due to environmental filtering caused by disturbances near roads influencing communities’ assemblies, causing changes in the communities’ functions, and phylogenies. Phylogenetic ecology brings the evolutionary history to the ecosystem functioning (Cavender-Bares et al., 2009; Ding et al., 2012; Srivastava et al., 2012; Webb et al., 2002) shedding light on the relation between ecological processes, selective pressures, and stability of an ecosystem (Cadotte et al., 2012; Helmus et al., 2010; Winter et al., 2013), when phenotypic differences and similarities among species are linked to evolutionary history (Webb, 2000). This demands the calculation of the phylogenetic signal of functional traits allowing inferences about niche conservatism and about convergences (Losos, 2008; Yang et al., 2014), as the link between phylogenetic diversity and functional diversity depends on the environmental filtering of traits that are conserved (i.e., homologous traits) or convergent (i.e., homoplasic traits) among phylogenetic lineages, causing the phylogenetic effects of clustering, and overdispersion, respectively (Cadotte et al., 2008; Cadotte and Davies, 2016). This relation between functions and phylogenies are especially meaningful in the Caatinga where plant functional traits determine resilience and resistance against pervasive anthropogenic disturbances (Carrión et al., 2017).

A great change is expected in functioning, phylogenetic structure, and diversity when ecosystems of severe environments, such as the Caatinga, are disturbed. As disturbances filter out some species, the filtered in species may increase in abundance meanwhile the community may lose some of its phylogenetic lineages (Helmus et al., 2007). Few studies have evaluated the effect of anthropogenic disturbances and resilience in the Caatinga (Albuquerque et al., 2012; Ribeiro et al., 2016, 2015), and even fewer have addressed the functional and phylogenetic structure of plant communities as effects of disturbances in general (Ding et al., 2012) and herbivory in particular (Carrión et al., 2017; Ribeiro et al., 2016). Species loss and phylogenetic clustering caused by disturbances may constrain the functioning, functional diversity and functional redundancy of Caatinga as a consequence of a decreased number of niches (Carrión et al., 2017). As far as we know, there have been no studies that have either assessed the functional and the phylogenetic effects of disturbances associated with roads in the Caatinga.

We aimed to evaluate the functional and the phylogenetic effects of disturbance near roads in the Caatinga. For that, we sampled plots near roads, and further from roads in order to measure differences concerning taxonomic, functional and phylogenetic diversities as well as phylogenetic structure and phylogenetic signal for disturbance-related traits. We tested the following hypotheses: (i) since roads are conduits of disturbances that filters out species, Caatinga near roads will exhibit lower taxonomic, functional and phylogenetic diversity compared to Caatinga further from roads; (ii) Caatinga near roads are more phylogenetically and functionally clustered than Caatinga further from roads; (iii) traits associated with herbivory deterrence are predominantly conserved within phylogenetic lineages of the Caatinga flora; and (iv) disturbance near roads alter the Caatinga functioning because of selection for disturbance-related traits.

## Materials and methods

### Study area

Our study was conducted in areas of Caatinga vegetation in northern Bahia State, Brazil. Occupying an area of 734,478 km^2^ (Silva et al., 2004), Caatinga vegetation is part of the global metacommunity of seasonally dry tropical forests (STDF) (Pennington et al., 2009) and represents the largest STDF in the Neotropics (Queiroz et al., 2017). The study area covers arboreal-shrub Caatinga or woody Caatinga of the Great Landscape Units of the Sertaneja Depression. The sampling area includes the roadsides of BR-235 from the municipality of Juazeiro to that of Jeremoabo. Roadsides in Brazil are poorly fenced, except for the main highways that are well-fenced. The roadside of the study area was completely unfenced before and during the sampling. The chosen private areas further from roads were completely unfenced as well. Domestic animals had free access to all studied areas. So, all studied area is grazed in some extent. The private areas are located a few kilometers distant from BR-235 inside a set of properties in the municipality of Canudos. The study site was chosen because of the roadsides with remaining Caatinga vegetation and because of the unfragmented and unfenced Caatinga from the roadsides up to the plots far from the road (i.e., up to 8,5 km from the road). Thus, the chosen study area presented a very specific situation of free movement of domestic animals, from the road to the plots, in an uninterrupted manner, and all plots were grazed by domestic animals to some extent, especially by goats that were constantly seen during the sampling period (Figure S1, S2, and S3).

The study area is located between 9°31’ and 10°05’S, and between 38°16’ and 40°07’W, and comprises four municipalities: Jeremoabo, Canudos, Curaçá and Juazeiro (Figures S1, and S3). According to the Köppen classification, the regional climate is BSh, typically semiarid, with mean annual rainfall lower than 550 mm, and elevations between 320-m and 520-m above sea level (Alvares et al., 2013; Carrión et al., 2017). The annual precipitation pattern is irregular, characterized by years of extreme drought, followed by an occasional year of torrential rains. Mean temperature varies between 23 to 27 °C, mean relative humidity is around 50% and the evapotranspiration rate is high (Daily and Mitchell, 2000). Despite the physiognomic variation of the Caatinga, the vegetation of all plots is patchy with clumps of woody shrubs and trees with a herb layer beneath (see Carrión et al., 2017).

### Sampling

We sampled all plant life forms in eight plots of 20-m x 50-m (1,000 m^2^) subdivided into subplots of 10-m x10-m (Figures S1, and S2). Four plots were allocated in different situations of Caatinga near road at distances from 30-m to 170-m from BR-235, and four plots in Caatinga far from roads at distances from 4,500-m to 8,500-m from BR-235, thus totaling eight plots and 0.8 ha (Figure S3). The distances were Log_10_ transformed for statistical analysis that used distances as continuous variable. We divided species into two components: woody (arboreal-shrubby) and non-woody (herbaceous-subshrub). All woody individuals with a circumference at ground level ≥ 10 cm and height ≥ 1 m, including woody lianas, were considered as the woody component of vegetation. The survey of non-woody species was carried out in two 10-m x 10-m subplots within each 20-m x 50-m plot (Figure S2), these being the first and last sampled subplots of the woody component (Figure S1). The cover of all non-woody species was estimated using the Braun-Blanquet method. The first five values (5 - 75 to 100% coverage, 4 -50 to 75%, 3 - 25 to 50%, 2 - 5 to 25% and 1 - 1 to 5%) of the Braun-Blanquet method refer to coverage while the last two value scales (+ - species with many individuals and low coverage, and r - rare species with low coverage) are primarily estimated by abundance (Munhoz and Araújo, 2011). From the Braun-Blanquet scale, we calculated coverage area (CA) and relative coverage (RC) using averaged values for each category (Kent, 2012).

We classified all species according to their life form using Raunkiaer’s life form system (Martins and Batalha, 2011). Species were also classified according to functional characteristics related to resilience/resistance to disturbances including resprouting ability, urticancy/toxicity and spinescent succulents, and according to functional characteristics affected by goat herbivory, including endozoochory, because of the fleshy fruits, and nitrogen fixation, because of the low C/N (Carrión et al., 2017; Leal et al., 2003; Severson and DeBano, 1991). Because of its dubious life form, the hemiepiphyte *Hylocereus setaceus* (Salm-Dyck) Ralf Bauer was considered in both, the woody and the non-woody community.

All of the species found with regrowth after a loss of aerial parts were considered resprouters, while all species with succulent and spiny structures were considered succulent spinescent (Carrión et al., 2017). Urticancy/toxicity was considered if species exhibited some vegetative organ with urticancy or that produce some toxic compound that affects goats, sheep and cattle (Bezerra et al., 2012). We considered nitrogen fixers as those species and genera that have a record of positive nodulation by nitrogen-fixing bacteria in the literature (Sprent, 2009). Species were considered endozoochoric through records in the literature or morphological observation (Pijl, 1982). In this study, secondary dispersion records were not considered.

In addition, we used maximum diameter and maximum height per species for the species of the woody component as quantitative functional traits. For this we calculated the mean for the three highest individuals and the three largest diameters of each species. We also used relative coverage (RC, in m^2^), calculated from Braun-Blanquet categories, of each non-woody species as a functional trait.

Species were recorded and later identified by expert consultation, herbaria comparison and/or specialized literature. The taxonomic system adopted was APG IV (The Angiosperm Phylogeny Group, 2016). We used information available on the Tropicos platform (Missouri Botanical Garden, 2018) and Taxonomic Name Resolution Service tool version 4.0 (Boyle et al., 2013) for appropriate species nomenclature and respective abbreviations of the authors.

### Statistical analyses and data

#### Road distance

We evaluated the effect of the logarithmic distance (Log_10_) from the road on total plant species richness, life form richness (Raunkiaer, 1934), endozoochorous richness (Pijl, 1982), resprouter richness (Meira-Neto et al., 2011) and nitrogen-fixer richness (Sprent, 2009). We performed generalized linear models (Glm) with Quasi-Poisson distributions for regressions associating and testing the significance of road distances with the different measures of species richness. The quasi-Poisson distributions were chosen when the dispersion parameters for gaussian family were too large. All analyses were performed in R statistical environment (R Development Core Team, 2015).

#### Functional traits

We evaluated differences in species richness and number of individuals of each functional trait between plots near roads, and plots further from roads. For these data, we used generalized linear models (Glm), Poisson family, with a logarithmic link function. When residual deviations were greater than the residual degrees of freedom, the standard errors were corrected using a Quasi-Poisson. We assessed significance using the Chi-squared test. All analyses were performed in R statistical environment (R Development Core Team, 2015).

#### Functional and phylogenetic diversity

Functional and phylogenetic diversities are different dimensions of the biodiversity and fundamental to test the hypothesis that Caatinga vegetation near roads will exhibit decreased taxonomic, functional and phylogenetic diversity compared to Caatinga vegetation far from roads. In order to analyze functional diversity, four indexes were used: functional richness (FRic), functional evenness (FEve), functional divergence (FDiv) (Villéger et al., 2008) and functional dispersion (FDis) (Laliberté and Legendre, 2010). Functional richness represents the amount of functional space filled by the community, while functional evenness quantifies the regularity that the functional space is filled by the species, weighing their abundances. Functional divergence can be understood as the distribution of the abundance occupied by the species within the volume of the functional space so that the divergence is low when the most abundant species is close to the center of the amplitude of the functional traits and high when this species has extreme trait values. Finally, functional dispersion refers to the average distance of one species in the multidimensional functional space for the centroid of all species.

We calculated these indices for all species sampled considering the characteristics related to tolerance/resistance strategies for herbivory deterrence: resprouting abilities, succulence with spines, urticancy/toxicity and annuals. For the woody component, calculations were made considering the following functional traits: resprouting abilitiy, urticancy/toxicity, succulents, nitrogen fixers, life form, endozoochory, maximum diameter (cm) and maximum height (m). For the non-woody component (herbaceous-sub-shrub), the functional traits considered were resprouting ability, annuals (therophytes), urticancy/toxicity, succulents, nitrogen fixers, life form and relative coverage of each species. All functional diversity indexes were calculated in R statistical environment (R Development Core Team, 2015), using the methods, scripts (Villéger et al., 2008) and the functions of the FD package (Laliberté and Legendre, 2010).

Phylogenetic diversity was calculated by the sum of the phylogenetic distance between species in each community using the PD (phylogenetic diversity) index provided by the Phylocom 4.2 program (Webb et al., 2011) and its standardized measure (standardized effect size - SESPD) calculated through ‘picante’ package (Kembel, 2015) in R statistical environment (R Development Core Team, 2015). The phylogeny used was constructed in the same way as the phylogenetic structure described in the next section.

To investigate the effect of disturbance near roads on variation in community functional and phylogenetic diversity, we tested the calculated indices statistically. We used Shapiro-Wilk test to verify normality of the data. We performed an ANOVA for the data with normal distribution and the Kruskal-Wallis test for distribution-free data (Zar, 1996).

#### Phylogenetic and functional structure

The phylogenetic and functional structure measures were taken to test the hypothesis that Caatinga vegetation near roads will be more phylogenetically and functionally clustered than Caatinga vegetation far from roads. For the phylogeny of the woody component, we used all the tree, shrub and liana species of the phytosociological survey identified at least to the family level (87 species). The phylogeny of herbaceous and sub-shrub species (non-woody component) included all non-woody species identified at least to the family level for a total of 86 species. Except for *Selaginella convoluta* (see phylogenetic signal section), all the morphospecies sampled in this survey were inserted into mega-tree R20160415.new (Gastauer and Meira-Neto, 2017) via phylomatic function, which was dated (Bell et al., 2010) using the algorithm bladj in combination with the file ‘ages’ in Phylocom (Gastauer and Meira-Neto, 2017; Webb et al., 2011). That mega- tree was chosen because it is an updated mega-tree using the updated APG IV classification (The Angiosperm Phylogeny Group, 2016).

With incidence data, we calculated phylogenetic structure for plots near roads and for plots further from roads, considering woody species or non-woody species, using the following phylogenetic measures with Phylocom 4.2 software (Webb et al., 2011): mean pairwise distance (MPD), mean nearest taxon distance (MNTD), net relatedness index (NRI) and nearest taxon index (NTI). Phylogenetic analysis was done using the unconstrained model of Phylocom (Gastauer and Meira-Neto, 2015; Webb et al., 2011) that maintains the species richness of each plot, but all species of the metacommunity (pool of species) have the same chance of being included (Webb et al., 2002) within each of the 10,000 randomizations. We defined the metacommunity as all the species sampled in the collection area (8,000m^2^), identified at species, genus or family level. We used the ‘comstruct’ function to calculate the phylogenetic metrics. We calculated these measures at different spatial scales: 10-m x 10-m and 20-m x 50-m.

In order to evaluate the functional structure of communities, we computed Gower distances among species based on the trait matrix (see above) using the ‘gowdis’ command from ‘FD’ package in R to perform the rarefaction and extrapolation of functional diversity (Villéger et al. 2008). From this we built a functional dendrogram in Newick format using ‘hclust2phylog’ from the ‘ade4’ package in R (Dray et al., 2018). We then computed traitNRI and traitNTI values for each plot using “ses.mpd” and “ses.mntd” commands and the unconstrained null model that assumes the same probability for all species to enter randomized communities. This works similar to the computation of phylogenetic community structure, but instead of phylogenetic distances among species of a plot, functional distances among species with the focal trait are calculated. The functional tree was built with all woody and non-woody species (n=179).

We tested statistical significance through a bilateral t-test for one sample, using the generated NRI and NTI metrics (Gastauer and Meira-Neto, 2015).

#### Phylogenetic signal

Phylogenetic signal was tested using two methods: the ‘aotf’ function in Phylocom 4.2 (Webb et al., 2011) for the quantitative traits (maximum diameter, maximum height and relative coverage), and calculating the traitNRI for the qualitative traits (Gastauer et al., 2017), except for *Selaginella convoluta*, which was not considered in the tests of the phylogenetic signal so as not to create bias caused by imbalanced phylogenetic trees because of the large phylogenetic distance (566 myr) between this only pteridophyte species and all the others (see Duchêne et al., 2015; Holman, 2005). For the endozoochory trait, maximum diameter and maximum height only the species of the woody component were considered (n=87). For the relative coverage trait (RC) only the species of the non-woody component were considered.

The ‘aotf’ module compares the variance rank of the observed mean variance of the trait of interest across all nodes from a null generated distribution of 9999 random repeats to trait values through phylogeny (Webb et al., 2011).

In order to evaluate trait conservatism, the significance of the variance of the independent contrasts was analyzed. 9999 randomizations were made and the trait was considered conserved if the variation of contrasts observed was less than 2.5% of the randomizations. TraitNRI and traitNTI were used to assert that all species possessing a given qualitative trait are more related to each other than expected by chance. The traitNRI was calculated for the phylogenetic structure of the community formed by the species with each trait, in order to emulate the phylogenetic signal of these traits. One thousand randomizations were performed and evaluated through null model. If traitNRIs or traitNTIs were lower than – 1.96 or higher than 1.96 (p<0.05), they were considered significant (Gastauer et al., 2017).

## Results

### Species richness

We recorded 178 species, of which seven angiosperms could not be identified, leaving 171 species identified for phylogenetic purposes: 86 woody species and 86 non-woody species (Table S1). We recorded 106 species near roads and 115 species in plots far from road. In plots near roads we sampled 1059 individuals belonging to 49 species, and in plots far from roads we sampled 1176 individuals belonging to 63 species. Plots near roads presented significantly lower species richness than plots far from roads (p<0.001). We sampled 57 non-woody species in plots near roads and 52 in plots far from roads. Species richness of non-woody plants did not differ between plots near roads and plots far from the roads (p=0.760).

### Road distance

Distance from the road into Caatinga was positively related to total species richness of plants, to phanerophyte richness, to resprouter richness and endozoochore richness, indicating overall species richness loss with the proximity of the road (Figure 1). The results show that disturbance decreases as the distance from the road increases. The species richness of chamaephytes, lianas and nitrogen-fixers did not present significant trends. The life form that experienced the greatest decrease in species richness with road-associated disturbance was phanerophytes while the functional group that experienced the greatest decrease in species richness with road-associated disturbance was the endozoochorous species-group.

**Figure 1:**
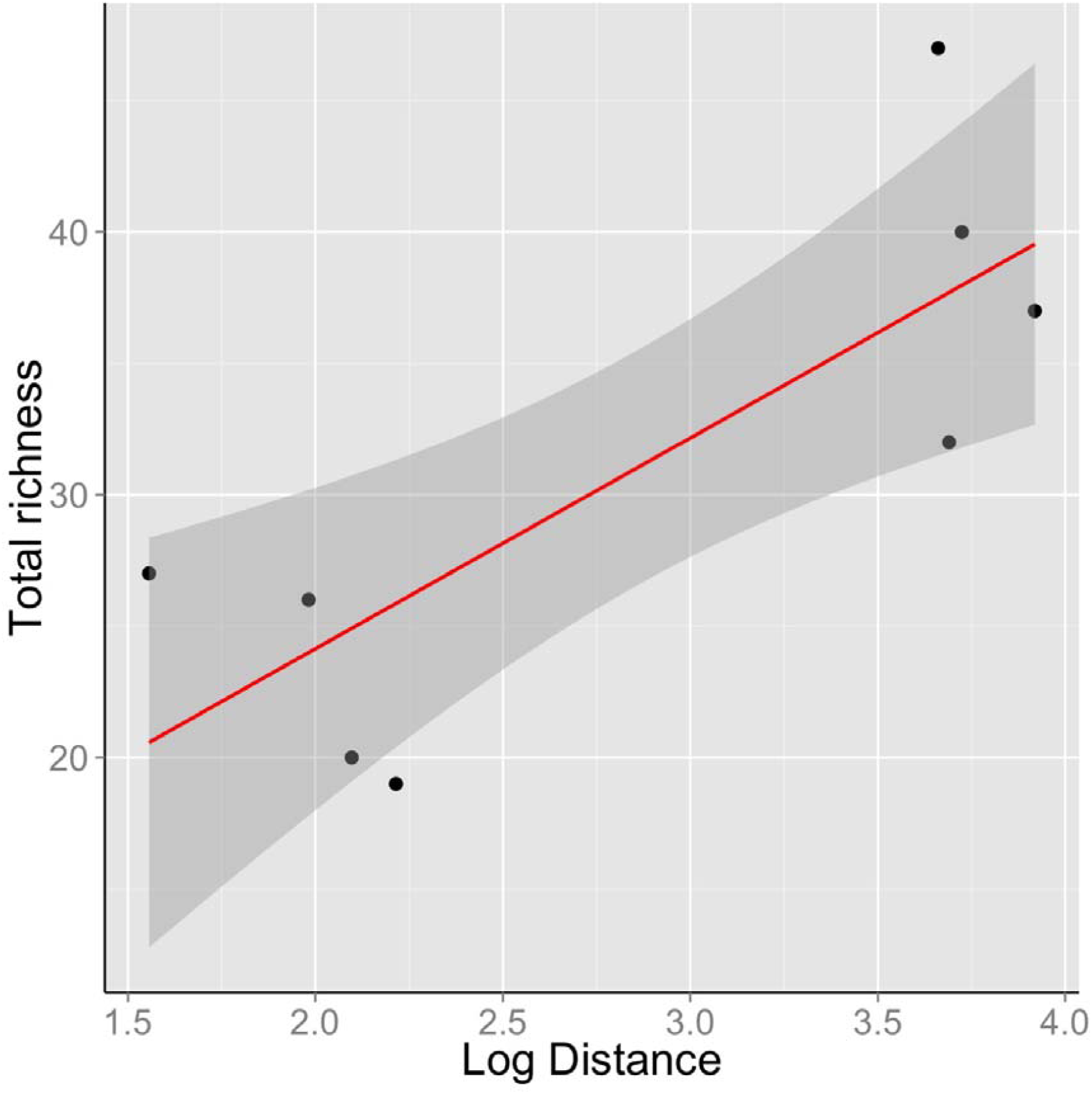
Generalized linear model of total plant species richness (Total richness) responding to Log Distance from the road (Log_10_ distance in meters); p<0.0181.

### Functional traits

Among woody species, 78 species were phanerophytes (n=2176), nine were woody lianas (n=48), one was a geophyte (n=10) and one was a hemiepiphyte (n=1). Among non-woody species 53 species were therophytes, 24 chamaephytes, six were hemicryptophytes, four were non-woody lianas, two were geophytes, one was an epiphyte and one was a hemiepiphyte. Plots far from roads had higher species richness of phanerophytes (p=0.0077) and of chamaephytes (p=0.047, Figure 2). In the woody component (trees, shrubs and lianas), 39 species were resprouters, 27 were endozoochoric, 16 were nitrogen-fixers, nine were urticant or toxic and seven were succulents with spines. Among the non-woody species, eight were spiny succulents, eight were nitrogen fixers, five were endozoochoric, three were resprouters and three were urticant or toxic (Table S1). The species richness of woody resprouters (p=0.027) as well as species richness of all resprouters (p=0.021) were lower near roads than far from roads (Figure 3). There were less species of woody nitrogen-fixers (p=0.018) as well as less species of nitrogen-fixers in general (p=0.027) in plots near roads than in plots far from roads. There were also less endozoochorous species (p=0.002) in plots near roads than in plots far from roads (Figure 3).

**Figure 2:**
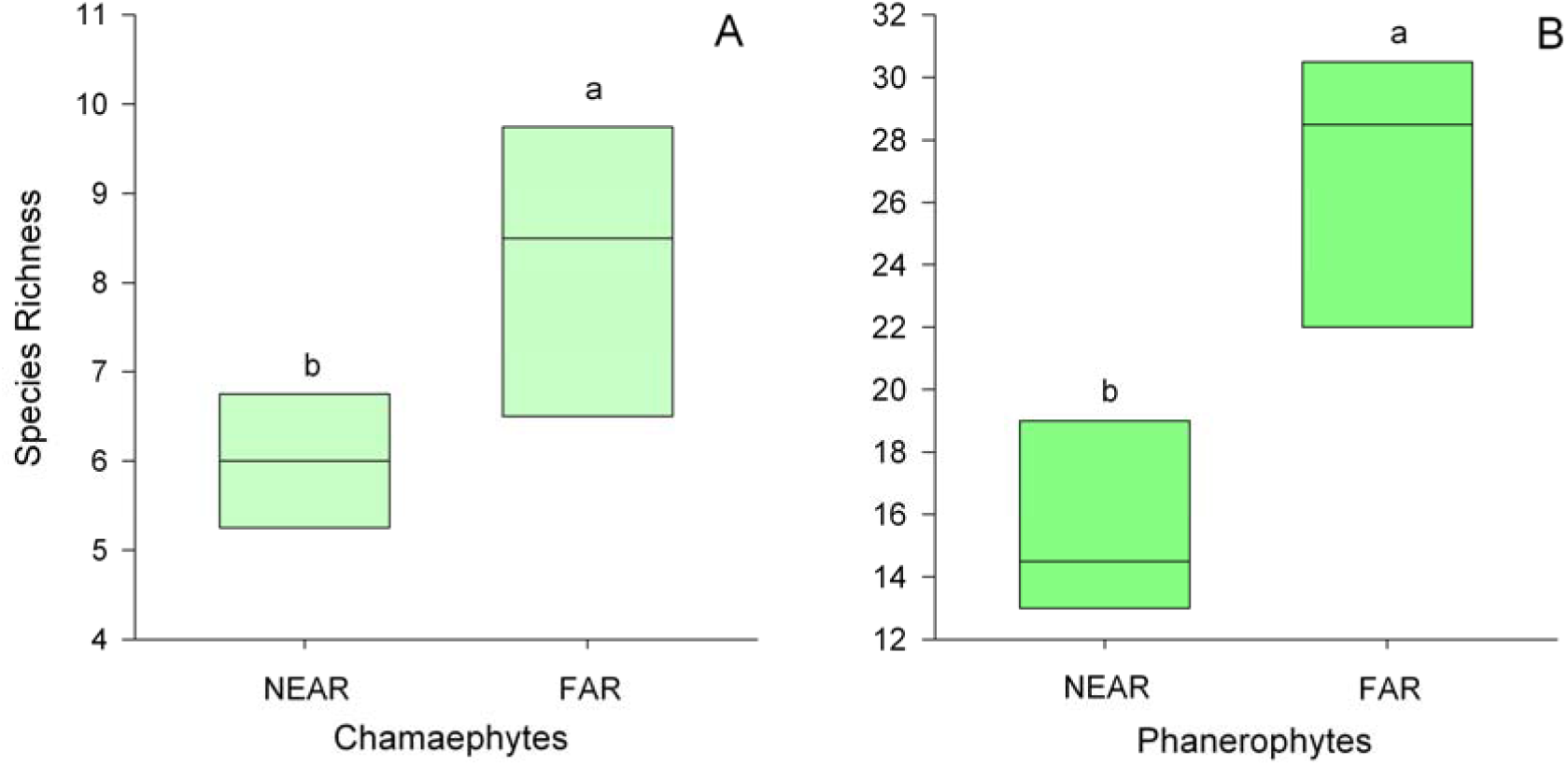
Boxplots of the richness of woody species and all species per 20 x 50-m plot per life form near the road (NEAR) and far from the road (FAR). GLM significance: A - p=0.0489; B - p=0.0077.

**Figure 3:**
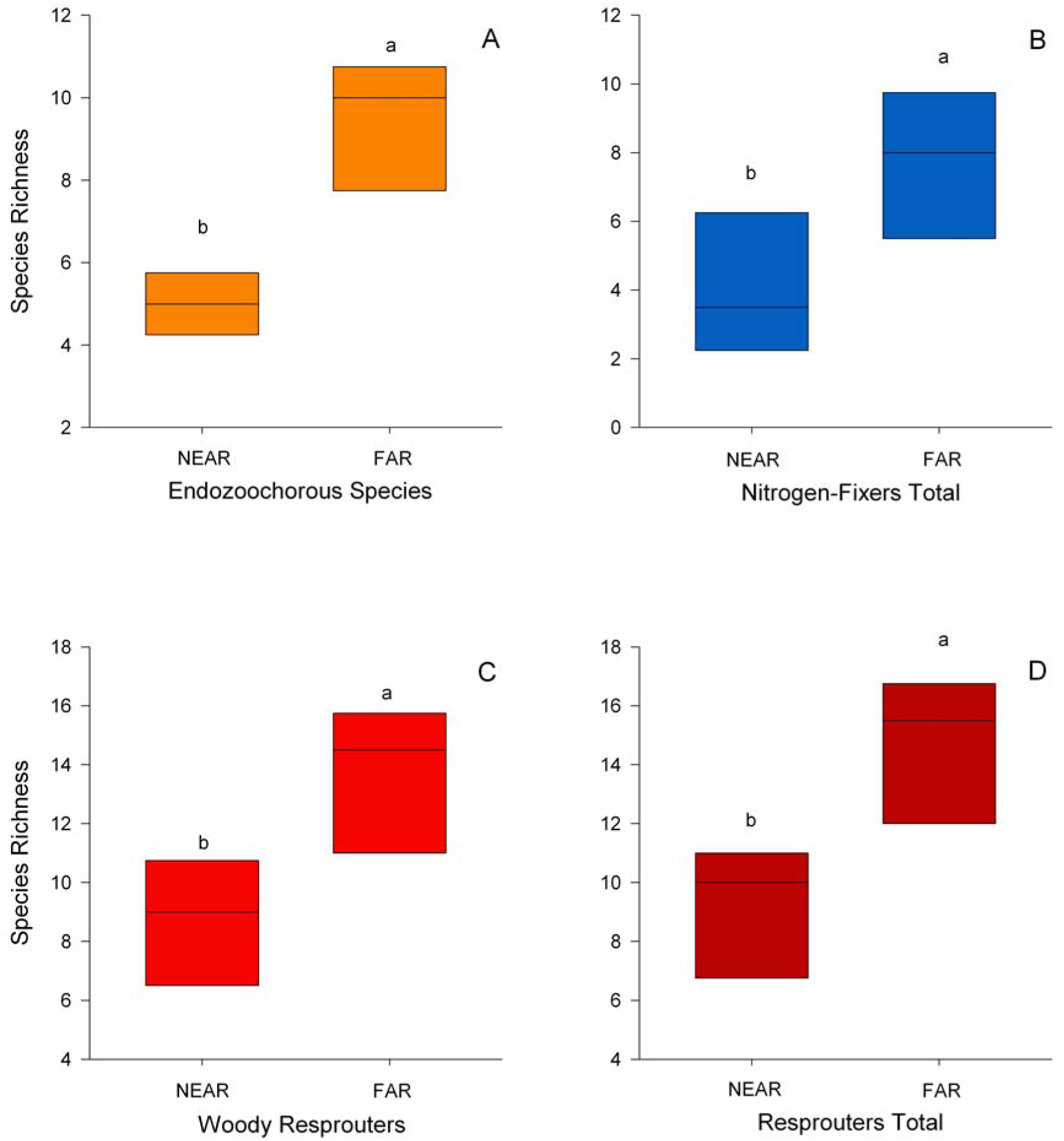
Boxplots of the richness of woody species and all species per 20 x 50-m plot of the herbivory-related functional traits (resprouters, endozoochorous and nitrogen fixers) near the road (NEAR) and far from the road (FAR). GLM significance for quasipoisson family distributions: A - p=0.0022; B - p=0.05; C – p=0.0286; D – p=0.0228).

### Functional and phylogenetic diversity

Among woody species, plots near roads presented lower Fric (p=0.0004) and PD (p=0.023) than plots far from roads. The other functional diversity attributes (FDiv, FEve and FDis), as well as sesPD, did not differ significantly (Table 1). Among non-woody species, the functional diversity metrics did not differ between samples near and far from roads. PD and sesPD were not significantly different as well (Table 1) among non-woody species. Functional diversity, in general, did not differ (FRic, FDiv, FEve, FDis) between the plots near and far from roads (Table 1).

**Table 1:**
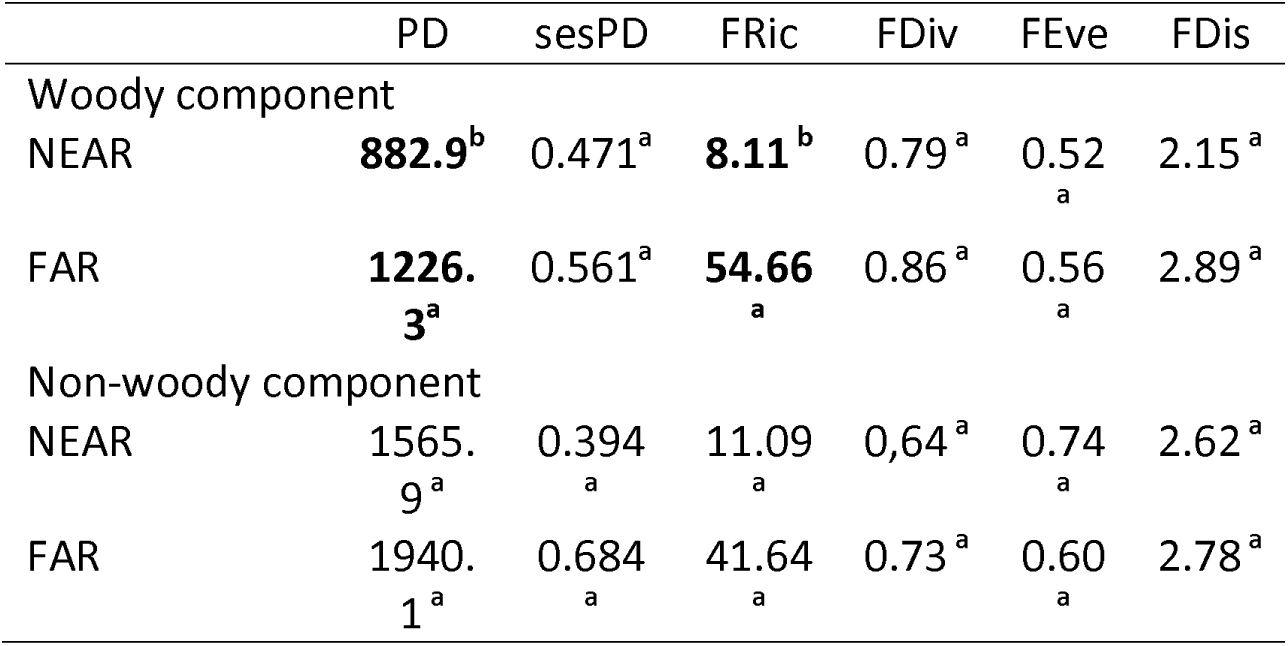
Phylogenetic diversity (PD and sesPD) and functional diversity (FRic - Functional Richness, FDiv - Functional Divergence, FEve - Functional Eveness and FDis - Functional Dispersion, for all functional attributes measured) of woody and non-woody species sampled in the Caatinga of northern Bahia, Brazil. Plots near roads (NEAR), plots far from roads (FAR). Bold values and different letters mean significant difference of an ANOVA for the data with normal distribution and the Kruskal-Wallis test for distribution-free data (p=0.05)

|  | PD | sesPD | FRic | FDiv | FEve | FDis |
| --- | --- | --- | --- | --- | --- | --- |
| Woody component |  |  |  |  |  |  |
| NEAR | **882.9<sup>b</sup>** | 0.471<sup>a</sup> | **8.11<sup>b</sup>** | 0.79<sup>a</sup> | 0.52<sub>a</sub> | 2.15<sup>a</sup> |
| FAR | **1226.3<sup>a</sup>** | 0.561<sup>a</sup> | **54.66<sub>a</sub>** | 0.86<sup>a</sup> | 0.56<sub>a</sub> | 2.89<sup>a</sup> |
| Non-woody component |  |  |  |  |  |  |
| NEAR | 1565.9<sup>a</sup> | 0.394<sub>a</sub> | 11.09<sub>a</sub> | 0.64<sup>a</sup> | 0.74<sub>a</sub> | 2.62<sup>a</sup> |
| FAR | 1940.1<sup>a</sup> | 0.684<sub>a</sub> | 41.64<sub>a</sub> | 0.73<sup>a</sup> | 0.60<sub>a</sub> | 2.78<sup>a</sup> |

### Phylogenetic and functional structure

The phylogenetic structure of Caatinga further from roads differed from randomness at the 10-m x 10-m scale (Table 2), with NRI significantly overdispersed for woody species (p=0.027) as well as for non-woody species (p=0.002). Those woody and non-woody species were phylogenetically overdispersed in plots further from roads for NRI (Table 2). Woody and non-woody species were phylogenetically clustered in plots near roads for NTI (Table 2). At the scale of 10-m x 10-m, the samples near roads presented NTI significantly clustered (Table 2) for woody species (p=0.037) and for non-woody species (p=0.031). Woody and non-woody species presented different phylogenetic structure at the 20-m x 50-m scale as well as at the 10-m x 10-m scale depending on the distance from roads. Further from roads, NRI shows phylogenetic overdispersion at both scales (Table 2, Table S3, Figure S2), whereas NTI shows phylogenetic clustering for plots near roads at both scales (Table 2).

**Table 2:** Net Relatedness Index (NRI) and Nearest Taxon Index (NTI) based on the unconstraint null model for the woody and non-woody component for the scale of 10-m x 10-m. Significant p- value indicates that the phylogenetic structure differs from zero according to the t-test for one sample. N=number of subplots of 10-m x 10-m. Plots near road (NEAR), and plots further from road (FAR). Bold values mean significance

|  |  | NRI |  |  | NTI |  |  |
| --- | --- | --- | --- | --- | --- | --- | --- |
|  | N | Mean | sd | p | Mean | sd | p |
| Woody component |  |  |  |  |  |  |  |
| NEAR | 40 | 0.092 | 0.87 | 0.502 | **0.335** | **0.98** | **0.037** |
| FAR | 40 | **-0.225** | **0.62** | **0.027** | 0.029 | 0.85 | 0.829 |
| Non-woody component |  |  |  |  |  |  |  |
| NEAR | 8 | 0.118 | 0.97 | 0.742 | **1.039** | **1.09** | **0.031** |
| FAR | 8 | **-0.636** | **0.38** | **0.002** | -0.696 | 0.89 | 0.061 |

Calculating phylogenetic structure only with plants possessing traits related to herbivory (resprouters, urticants or toxic, succulents with spines), the traitNTI was significantly higher than zero in plots near roads, indicating functional clustering and closer relatedness among individuals than by chance (Table 3). Calculating phylogenetic structure with plants with all traits, the traitNRI of plots near roads was significantly greater than zero, indicating phylogenetic clustering, that is, closer relatedness than by chance (Table 3). These results show that environmental filtering near roads filters in species resilient/resistant to herbivory throughout the entire phylogenetic tree, but especially towards the tips of the phylogenetic branches.

**Table 3:**
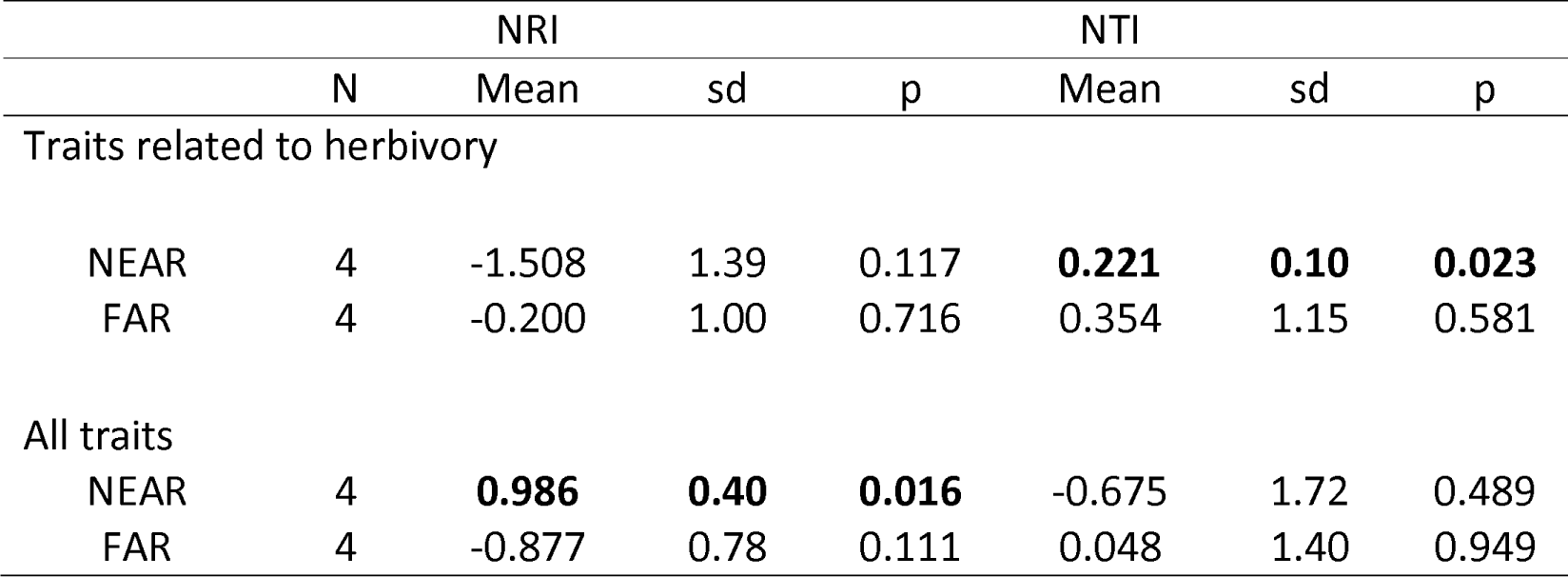
Net Relatedness Index (NRI) and Nearest Taxon Index (NTI) of the functional structure for traits related to herbivory and for all traits. Significant p-value indicates that the phylogenetic structure differs from zero according to the t-test for one sample. N=number of plots (20-m x 50- m). Plots near road (NEAR), and plots further from road (FAR). Bold values mean significance

### Phylogenetic signal

We did not find phylogenetic signal for maximum diameter (contrast variation=6.808, p>0.05) and maximum height (contrast variation=0.165; p>0.05) as well as herbaceous relative cover (contrast variation=0.043, p>0.05). Since from comstruct output it becomes possible to compute significance levels for phylogenetic structure values to be different from the zero expectation (MPD.rankHigh/nruns), Nitrogen fixers were clustered phylogenetically, (NRI=20.97), as well as those with resprouting ability (NRI=4.43), with urticancy or toxicity (NRI=9.54), spiny succulents (NRI=9.55), phanerophytes (NRI=5.70) and lianas (NRI=3.22), demonstrating that those functional traits related to herbivory and long-stemmed life forms are conserved within phylogenetic lineages of the metacommunity tree. In contrast, the endozoochorous species (NRI=- 1.44), chamaephytes (NRI=0.39), annuals/therophytes (NRI=0.51) and hemicryptophytes (NRI=- 0.29) were not significantly clustered.

Regarding the traits related to strategies for herbivory deterrence, distribution of traits on the phylogenetic tree of the woody component shows spinescent succulents restricted to the families Cactaceae and Bromeliaceae. Meanwhile, resprouters were distributed mainly among the Rosids clade. Urticancy and toxicity was mainly distributed in the family Euphorbiaceae, with some representatives in the families Fabaceae, Apocynaceae and Portulacaceae. Annuals were distributed throughout the metacommunity phylogenetic tree, but mainly in the families Poaceae, Commelinaceae, Scrophulariaceae, Asteraceae, Amaranthaceae and Cleomaceae (Figure 4).

**Figure 4:**
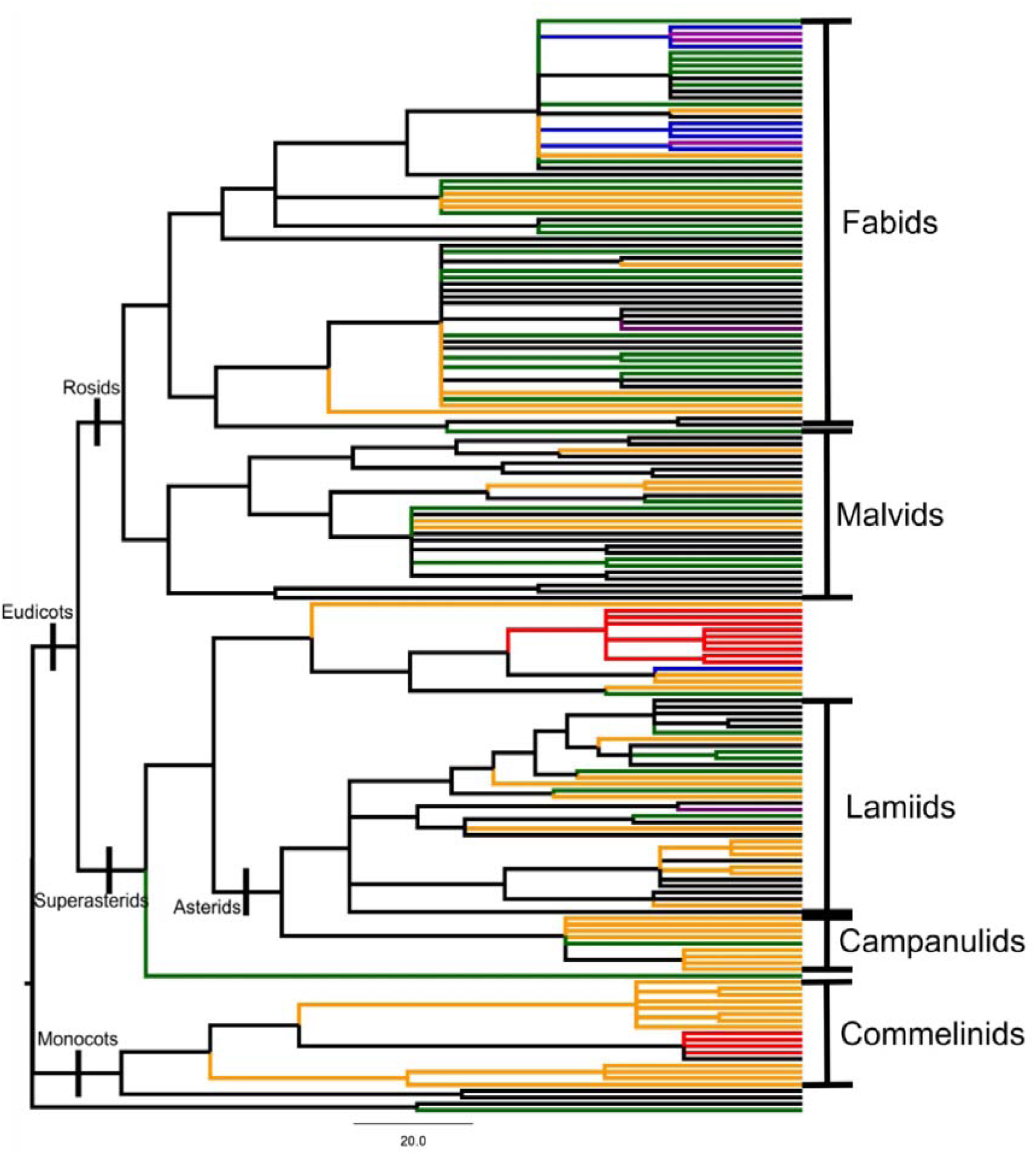
Phylogenetic tree of all species. The marked branches highlight four functional traits: resprouters - green; urticancy and/or toxic species - blue; succulent species with spines - red; annuals (therophytes) - orange; urticancy and/or toxic resprouters – purple (adapted from Carrión et al., 2017).

## Discussion

Plant species richness decreased in plots near roads, suggesting roads as sources, and/or conduits of anthropogenic disturbance in the Caatinga. Plots near roads showed decreased taxonomic, functional and phylogenetic diversity compared to plots far from roads. The phylogenetic structure of Caatinga near roads is clustered, indicating road-associated disturbances as environmental filters since traits associated to livestock herbivory deterrence were predominantly conserved within phylogenetic lineages and preferentially selected near roads. Therefore, roads should be considered source, and/or conduits of disturbances causing an impoverishment of the biodiversity, and the ecosystem functioning in the Caatinga.

Plots near roads presented lower species richness than plots further from roads. There are fewer species of chamaephytes, phanerophytes, endozoochorous, nitrogen-fixers and resprouters in plots near roads than plots further from roads, thus, there was a loss of redundancy for these functional groups. The communities near roads were phylogenetically more closely related because of some recent lineages filtered in by the influence of the road-associated disturbances, according to the NTI of woody and non-woody communities. Overall species richness, phanerophyte species richness, resprouter species richness and endozoochorous species richness increased as the distance from the road increased. These results suggest that roads are sources, and/or conduits of disturbances (Forman and Alexander, 1998; Forman and Deblinger, 2000) in Caatinga. Therefore, the results show that road vicinity holds environmental filters that cause species loss in different functional groups found in Caatinga.

Plots further from roads presented greater phylogenetic diversity and functional richness of woody species than plots near roads. Plots further from roads were phylogenetically overdispersed among woody, and non-woody species, congruently to the phylogenetically clustered plots near roads. Since the set of traits related to herbivory have a phylogenetic signal (i.e. resprouter ability, succulence with spines, and urticancy/toxicity), this effect suggests environmental filtering. Plots near roads were phylogenetically clustered throughout the entire phylogenetic tree (NRI) concerning the whole set of functional traits. Therefore, the phylogenetic clustering of plots close to the roads and the phylogenetic overdispersion further from the roads indicate changes in the functional structure of the Caatinga and show a decline in species richness in some parts of the functional space (Vamosi et al., 2009), as detected for chamaephytes, phanerophytes, endozoochores, lianas, nitrogen-fixers and resprouters. Thus, the working hypotheses i) that Caatinga near roads will exhibit lower taxonomic, functional and phylogenetic diversity compared to Caatinga further from roads, and ii) that Caatinga near roads will be more phylogenetically and functionally clustered than Caatinga further from roads, our results show that both hypotheses were confirmed. Our results also show that Caatinga communities near roads are more phylogenetically and functionally clustered among species with traits related to herbivory than communities further from roads. Consequently, disturbance amplified near roads cause loss of phylogenetic lineages across different life forms, different functional groups, causing evolutionary history loss. These results mean that disturbance is an old pressure (i.e. traitNRI for traits related to all traits), but herbivory is rather a selective pressure towards the tip of the phylogenetic tree of the Caatinga metacommunity (i.e. traitNTI for traits related to herbivory).

As a disturbance may select associated functional traits (Ding et al., 2012; Helmus et al., 2010), in the Caatinga near roads, functional traits related to herbivory deterrence (Carrión et al., 2017) should be selected. Since herbivory-deterrence traits are clustered, frequent disturbances may cause phylogenetic clustering by filtering in species with herbivory-deterrence traits, meanwhile absence of disturbances would lead to another phylogenetic structure (Vamosi et al., 2009). Congruently, our results showed plots near roads to be functionally and phylogenetically clustered. Therefore, those results not only are congruent with the herbivory-deterrence traits conserved within phylogenetic lineages in Caatinga, but also show that disturbances near roads alter plant communities because of selection for herbivory-deterrence traits, confirming the hypothesis that iii) traits associated with herbivory deterrence are predominantly conserved within phylogenetic lineages of the Caatinga flora.

Bovines, goats and sheep are domestic animals that cause intense herbivory, but herbivory by goats is usually the most pervasive (Rainbolt and Coblentz, 1999). Goats seek thoroughly the most palatable species, thereby imposing selective disturbance on the Caatinga flora for the benefit of less palatable species such as *Mimosa* ssp., *Cenostigma* ssp. and *Croton* ssp., which become abundant in high disturbed Caatinga (Alves et al., 2008). Therefore, in plots near roads, some species are being replaced by other more resilient/resistant. We found decreased species richness in several functional groups of species in plots near roads (Figure 1). Our results showed decreased richness of species with endozoochory in plots near roads congruently with reports that goat herbivory might reduce the abundance and diversity of succulent fruit species (Leal et al., 2005). Furthermore, we found that 80% of the woody individuals sampled in plots near roads were resprouters, while in plots further from roads 53% of the individuals were resprouters. Thus, in more disturbed Caatinga communities, fewer species benefit and may eventually proliferate, especially those that are less palatable, have resprouting ability and are non-endozoochorous, reducing rangeland and support for dispersers. That reduced rangeland and support for dispersers superimpose with road avoidance due to traffic noise and to road construction (Forman and Alexander, 1998; Forman and Deblinger, 2000) with ecological impact for ecosystem functioning (Laurance et al., 2009), constraining fauna in general and dispersers in particular. Therefore, roads reduce the diversity and functioning of the Caatinga, meanwhile promote species resistant and resilient to disturbances.

As disturbances associated with roads are rather chronic than sporadic, a persistent and consistent diversity loss on many levels may generate a gradient of disturbance initiating from the road/roadsides and radiating further into the Caatinga. In the Caatinga, species loss due to chronic anthropogenic disturbances does not occur randomly or uniformly, but rather in clusters throughout the phylogenetic tree (Ribeiro et al., 2016), and our results show that the same effects are caused near roads. Therefore, roads might be understood as axes of high human disturbances into the Caatinga, and be used as a proxy for habitat degradation (Leal et al., 2014; Ribeiro et al., 2015). As a consequence, on a large scale, a road network may produce cells with inner areas less disturbed and with peripheral areas more disturbed bordered by roads. This patchwork-like landscape may determine not only isolation and fragmentation in the Caatinga (Santos and Tabarelli, 2002), but also much of the phylogenetic, functional and taxonomic patterns of extant Caatinga areas.

We did not find differences in non-woody species richness and non-woody functional diversity between plots near roads and further from roads. These results indicate that these non-woody communities are more resilient/resistant to disturbances than woody communities. As our plots comprise predominantly patchy Caatinga, a physiognomy caused by facilitation (Carrión et al., 2017), the non-woody species seem to be less affected by disturbances inside the clumps/patches, being nursed by copious branched *Cenostigma* trees, urticant/toxic leaves of Euphorbiaceae and spinescent Cactaceae. Therefore, disturbances from roads can be minimized by facilitation that must be consider during the planning of restoration and conservation actions near roads.

Our results show roads as source/conduits of disturbances that cause loss of taxonomic, phylogenetic and functional diversity among plants as well as constrain fauna in general and dispersers in particular, causing an impoverishment of the ecosystem functioning in the Caatinga. Biodiversity conservation, and restoration planning must avoid the influence of roads close to or within natural areas in the Caatinga, and where roads are close, restoration and conservation practices may consider to use facilitation in order to increase diversity and functions.

## Acknowledgments

The authors thank the DNIT (Departamento Nacional de Infraestrutura de Transportes, process 50600.056799/2013-21), FUNARBE (process 6991), and BR235/UFV Team (process 1071/2013-DPP) for grants and financial support. The authors also thank CNPq (301913/2012-9), CAPES and FAPEMIG (APQ-01309-16), PPGBot and PPGEco of Universidade Federal de Viçosa for providing infrastructure, grants and scholarships; Dra. Ricarda Riina (MA), Dra. Maria Iracema Bezerra Loiola (UFC), Dra. Maria Staf (PMA), Dr. Marcos Sobral (UFSJ), Dr. Luciano Paganucci de Queiroz and UEFS Herbarium for helping with plant identification; Biodiversitas Foundation, Caboclo and Marcos Vinicius Varjão Romão for fieldwork help; and Evandro Fortini for helping with the figures. JAAMN holds a productivity fellowship (CNPq 307591/2016-6).

