## Supplementary Tables S1, S2, and Figures S1, S2, and S3 for "Roads as conduits of taxonomic, functional and phylogenetic degradation in caatinga vegetation"

**Supplementary Material**

**Table S1:** List of species sampled in eight plots of Caatinga vegetation in northern Bahia. LF: life form; Dmax: maximum diameter (only for woody species); Hmax: maximum height (only for woody species); RC: relative coverage (only for non-woody species); Res: species with regrowth capacity (X); Urt/tox: urticancy and/or toxic species (X); Suc: spinescent succulent species (X); Fix: nitrogen fixing species (X); Annual: species with annual life cycle (X); Endzoo: primary endozoochorous species (X).

| **Family**/Species | LF | Dmax | Hmax | RC | Res | Urt/tox | Suc | Fix | Annual | Endzoo |
| --- | --- | --- | --- | --- | --- | --- | --- | --- | --- | --- |
| **Acanthaceae** |  |  |  |  |  |  |  |  |  |  |
| *Harpochilus neesianus* Mart. ex Nees | FAN | 22.71 | 2.67 | - | X | 0 | 0 | 0 | 0 | 0 |
| *Ruellia* sp. | TER | - | - | 0.01 | 0 | 0 | 0 | 0 | X | - |
| **Amaranthaceae** |  |  |  |  |  |  |  |  |  |  |
| *Alternanthera brasiliana* (L.) Kuntze | TER | - | - | 3.99 | 0 | 0 | 0 | 0 | X | - |
| **Anacardiaceae** |  |  |  |  |  |  |  |  |  |  |
| *Myracrodruon urundeuva* Allemão | FAN | 32.04 | 9.00 | - | 0 | 0 | 0 | 0 | 0 | 0 |
| *Spondias tuberosa* Arruda | FAN | 61.12 | 6.00 | - | 0 | 0 | 0 | 0 | 0 | X |
| **Annonaceae** |  |  |  |  |  |  |  |  |  |  |
| *Annona leptopetala* (R.E. Fr.) H. Rainer | FAN | 11.95 | 4.67 | - | 0 | 0 | 0 | 0 | 0 | X |
| *Annona spinescens* Mart. | FAN | 25.83 | 4.17 | - | X | 0 | 0 | 0 | 0 | X |
| **Apocynaceae** |  |  |  |  |  |  |  |  |  |  |
| Apocynaceae 1 | LIA | - | - | 0.01 | 0 | 0 | 0 | 0 | 0 | - |
| *Aspidosperma parvifolium* A. DC. | FAN | 24.77 | 6.17 | - | X | X | 0 | 0 | 0 | 0 |
| **Asteraceae** |  |  |  |  |  |  |  |  |  |  |
| *Conocliniopsis prasiifolia* R.M. King & H. Robinson | HEM | - | - | 3.63 | X | 0 | 0 | 0 | 0 | - |
| *Lepidaploa chalybaea* (Mart. ex DC.) H. Rob. | TER | - | - | 0.01 | 0 | 0 | 0 | 0 | X | - |
| *Lepidaploa cotoneaster* (Willd. ex Spreng.) H.Rob. | TER | - | - | 0.47 | 0 | 0 | 0 | 0 | X | - |
| *Lepidaploa* sp.1 | TER | - | - | 0.01 | 0 | 0 | 0 | 0 | X | - |
| *Lepidaploa* sp.2 | TER | - | - | 0.01 | 0 | 0 | 0 | 0 | X | - |
| Asteraceae 1 | TER | - | - | 2.32 | 0 | 0 | 0 | 0 | X | - |
| Asteraceae 2 | TER | - | - | 0.33 | 0 | 0 | 0 | 0 | X | - |
| Asteraceae 3 | TER | - | - | 0.33 | 0 | 0 | 0 | 0 | X | - |
| Asteraceae 4 | TER | - | - | 0.01 | 0 | 0 | 0 | 0 | X | - |
| **Bignoniaceae** |  |  |  |  |  |  |  |  |  |  |
| *Anemopaegma laeve* DC. | CAM | - | - | 0.01 | 0 | 0 | 0 | 0 | 0 | - |
| Bignoniaceae 1 | LIA | 10.44 | 6.30 | - | 0 | 0 | 0 | 0 | 0 | 0 |
| Bignoniaceae 2 | LIA | - | - | 0.01 | 0 | 0 | 0 | 0 | X | - |
| *Fridericia* sp.1 | LIA | 14.46 | 6.50 | - | 0 | 0 | 0 | 0 | 0 | 0 |
| *Fridericia* sp.2 | LIA | 8.11 | 2.17 | - | 0 | 0 | 0 | 0 | 0 | 0 |
| *Handroanthus spongiosus* (Rizzini) S.O. Grose | FAN | 45.78 | 7.33 | - | X | 0 | 0 | 0 | 0 | 0 |
| **Boraginaceae** |  |  |  |  |  |  |  |  |  |  |
| *Varronia globosa* Jacq. | FAN | 9.55 | 1.95 | - | 0 | 0 | 0 | 0 | 0 | X |
| **Bromeliaceae** |  |  |  |  |  |  |  |  |  |  |
| *Bromelia laciniosa* Mart. ex Schult. & Schult.f. | CAM | - | - | 5.61 | 0 | 0 | X | 0 | 0 | - |
| *Encholirium spectabile* Mart. ex Schult. & Schult.f. | HEM | - | - | 0.33 | 0 | 0 | X | 0 | 0 | - |
| *Hohenbergia catingae* Ule | HEM | - | - | 3.63 | 0 | 0 | X | 0 | 0 | - |
| *Neoglaziovia variegata* (Arruda) Mez | CAM | - | - | 7.79 | 0 | 0 | X | 0 | 0 | - |
| *Tillandsia recurvata*(L.) L. | EPI | - | - | 0.67 | 0 | 0 | 0 | 0 | 0 | - |
| **Burseraceae** |  |  |  |  |  |  |  |  |  |  |
| *Commiphora leptophloeos* (Mart.) J.B. Gillett | FAN | 66.74 | 7.33 | - | 0 | 0 | 0 | 0 | 0 | X |
| **Cactaceae** |  |  |  |  |  |  |  |  |  |  |
| *Cereus jamacaru* DC. | FAN | 21.65 | 5.17 | - | 0 | 0 | X | 0 | 0 | X |
| *Hylocereus setaceus* (Salm-Dyck ex DC.) Ralf Bauer | HEP | 3.18 | 1.20 | 1.18 | 0 | 0 | X | 0 | 0 | X |
| *Melocactus zehntneri* (Britton & Rose) Luetzelb. | CAM | - | - | 0.69 | 0 | 0 | X | 0 | 0 | - |
| *Pilosocereus catingicola* (Gürke) Byles & G.D. Rowley | FAN | 19.95 | 5.00 | - | 0 | 0 | X | 0 | 0 | X |
| *Pilosocereus gounellei* (F.A.C. Weber) Byles & G.D. Rowley | FAN | 33.68 | 2.50 | - | 0 | 0 | X | 0 | 0 | X |
| *Pilosocereus pachycladus* F. Ritter | FAN | 23.28 | 5.17 | - | 0 | 0 | X | 0 | 0 | X |
| *Pilosocereus tuberculatus* (Werderm.) Byles & G.D. Rowley | FAN | 9.39 | 2.83 | - | 0 | 0 | X | 0 | 0 | X |
| *Tacinga inamoena* (K.Schum.) N.P.Taylor & Stuppy | CAM | - | - | 2.66 | 0 | 0 | X | 0 | 0 | - |
| *Tacinga palmadora* (Britton & Rose) N.P. Taylor & Stuppy | FAN | 16.55 | 3.90 | - | 0 | 0 | X | 0 | 0 | X |
| **Capparaceae** |  |  |  |  |  |  |  |  |  |  |
| *Cynophalla flexuosa* (L.) J. Presl | LIA | 5.27 | 3.67 | - | 0 | 0 | 0 | 0 | 0 | X |
| *Neocalyptrocalyx longifolium* (Mart.) Cornejo & Iltis | FAN | 11.75 | 4.17 | - | X | 0 | 0 | 0 | 0 | X |
| **Cleomaceae** |  |  |  |  |  |  |  |  |  |  |
| *Cleome* sp. | TER | - | - | 0.03 | 0 | 0 | 0 | 0 | X | - |
| *Physostemon guianense* (Aubl.) Malme | TER | - | - | 0.02 | 0 | 0 | 0 | 0 | X | - |
| **Combretaceae** |  |  |  |  |  |  |  |  |  |  |
| *Combretum monetaria* Mart. | LIA | 4.45 | 4.50 | - | 0 | 0 | 0 | 0 | 0 | 0 |
| **Commelinaceae** |  |  |  |  |  |  |  |  |  |  |
| *Callisia filiformis* (M.Martens & Galeotti) D.R.Hunt | TER | - | - | 0.01 | 0 | 0 | 0 | 0 | X | - |
| *Commelina erecta* L. | TER | - | - | 0.70 | 0 | 0 | 0 | 0 | X | - |
| Commelinaceae 1 | TER | - | - | 0.01 | 0 | 0 | 0 | 0 | X | - |
| **Convolvulaceae** |  |  |  |  |  |  |  |  |  |  |
| *Evolvulus* sp. | TER | - | - | 0.48 | 0 | 0 | 0 | 0 | X | - |
| *Evolvulus* sp.2 | TER | - | - | 0.46 | 0 | 0 | 0 | 0 | X | - |
| *Evolvulus* sp.3 | TER | - | - | 0.47 | 0 | 0 | 0 | 0 | X | - |
| *Ipomoea brasiliana* Meisn. | GEO | 13.30 | 3.67 | - | 0 | 0 | 0 | 0 | 0 | 0 |
| *Jacquemontia* sp. | TER | - | - | 0.01 | 0 | 0 | 0 | 0 | X | - |
| *Jacquemontia* sp.2 | TER | - | - | 0.33 | 0 | 0 | 0 | 0 | X | - |
| *Merremia cissoides*(Lam.) Hallier f. | LIA | - | - | 0.01 | 0 | 0 | 0 | 0 | 0 | - |
| *Turbina cordata* (Choisy) D.F. Austin & Staples | LIA | 18.04 | 5.67 | - | 0 | 0 | 0 | 0 | 0 | 0 |
| **Cucurbitaceae** |  |  |  |  |  |  |  |  |  |  |
| *Apodanthera glaziovii* Cogn. | LIA | 3.50 | 3.00 | - | 0 | 0 | 0 | 0 | 0 | X |
| *Apodanthera trifoliata* Cogn. | LIA | 3.34 | 1.50 | - | 0 | 0 | 0 | 0 | 0 | X |
| **Dioscoreaceae** |  |  |  |  |  |  |  |  |  |  |
| *Dioscorea* sp. | GEO | - | - | 0.01 | 0 | 0 | 0 | 0 | 0 | - |
| *Dioscorea* sp.2 | GEO | - | - | 0.01 | 0 | 0 | 0 | 0 | 0 | - |
| **Erythroxylaceae** |  |  |  |  |  |  |  |  |  |  |
| *Erythroxylum* *betulaceum* Mart. | FAN | 7.34 | 2.23 | - | X | 0 | 0 | 0 | 0 | 0 |
| *Erythroxylum* *caatingae* Plowman | FAN | 12.41 | 2.50 | - | 0 | 0 | 0 | 0 | 0 | 0 |
| *Erythroxylum* *revolutum* Mart. | FAN | 9.53 | 3.93 | - | X | 0 | 0 | 0 | 0 | 0 |
| **Euphorbiaceae** |  |  |  |  |  |  |  |  |  |  |
| *Acalypha brasiliensis* Müll. Arg. | FAN | 17.93 | 2.83 | - | X | 0 | 0 | 0 | 0 | 0 |
| *Cnidoscolus loefgrenii* (Pax & K. Hoffm.) Pax & K. Hoffm. | HEM | - | - | 0.33 | 0 | X | 0 | 0 | 0 | - |
| *Cnidoscolus pubescens* Pohl | FAN | 32.27 | 6.67 | - | X | X | 0 | 0 | 0 | 0 |
| *Cnidoscolus quercifolius* Pohl | FAN | 10.61 | 5.00 | - | X | X | 0 | 0 | 0 | 0 |
| *Cnidoscolus* *urens* (L.) Arthur | CAM | - | - | 3.96 | 0 | X | 0 | 0 | 0 | - |
| *Croton arenosus* Carn.-Torres & Cordeiro | FAN | 6.41 | 1.57 | - | X | 0 | 0 | 0 | 0 | 0 |
| *Croton argyrophyllus* Kunth | FAN | 13.37 | 4.00 | - | X | 0 | 0 | 0 | 0 | 0 |
| *Croton blanchetianus* Baill. | FAN | 20.27 | 5.67 | - | X | 0 | 0 | 0 | 0 | 0 |
| *Croton echioides* Baill. | FAN | 18.99 | 8.17 | - | X | 0 | 0 | 0 | 0 | 0 |
| *Croton grewioides* Baill. | FAN | 12.94 | 2.77 | - | 0 | 0 | 0 | 0 | 0 | 0 |
| *Croton heliotropiifolius* Kunth | FAN | 3.50 | 1.20 | - | X | 0 | 0 | 0 | 0 | 0 |
| *Croton* sp. | CAM | - | - | 0.33 | 0 | 0 | 0 | 0 | 0 | - |
| *Croton virgultosus* Müll. Arg. | FAN | 8.91 | 2.00 | - | 0 | 0 | 0 | 0 | 0 | 0 |
| *Ditaxis desertorum* (Müll. Arg.) Pax & K. *Hoffm.* | FAN | 9.74 | 1.67 | - | X | 0 | 0 | 0 | 0 | 0 |
| *Euphorbia nutans* Lag. | TER | - | - | 0.66 | 0 | 0 | 0 | 0 | X | - |
| *Euphorbia* sp. | CAM | - | - | 0.01 | 0 | 0 | 0 | 0 | 0 | - |
| *Jatropha mollissima* (Pohl) Baill. | FAN | 6.21 | 4.23 | - | 0 | X | 0 | 0 | 0 | 0 |
| *Jatropha mutabilis* (Pohl) Baill. | FAN | 7.32 | 4.07 | - | 0 | X | 0 | 0 | 0 | 0 |
| *Jatropha ribifolia* (Pohl) Baill. | FAN | 3.18 | 1.50 | - | 0 | X | 0 | 0 | 0 | 0 |
| *Manihot* sp. | FAN | 8.54 | 3.93 | - | 0 | X | 0 | 0 | 0 | 0 |
| *Manihot* sp.2 | FAN | 10.16 | 5.17 | - | X | X | 0 | 0 | 0 | 0 |
| *Microstachys corniculata* (Vahl) Griseb. | TER | - | - | 0.34 | 0 | 0 | 0 | 0 | X | - |
| *Sapium glandulosum* (L.) Morong | FAN | 22.49 | 6.67 | - | X | 0 | 0 | 0 | 0 | 0 |
| *Stillingia trapezoidea* Ule | FAN | 4.19 | 2.13 | - | 0 | 0 | 0 | 0 | 0 | 0 |
| **Fabaceae** |  |  |  |  |  |  |  |  |  |  |
| *Aeschynomene martii* Benth. | FAN | 4.78 | 3.00 | - | 0 | 0 | 0 | X | 0 | 0 |
| *Calliandra depauperata* Benth. | FAN | 8.90 | 2.33 | - | X | 0 | 0 | X | 0 | 0 |
| *Chamaecrista repens* (Vogel) H.S.Irwin & Barneby | CAM | - | - | 0.01 | 0 | 0 | 0 | X | 0 | - |
| *Chamaecrista venulosa* (Benth.) H.S.Irwin & Barneby | TER | - | - | 0.33 | 0 | 0 | 0 | X | X | - |
| *Copaifera martii* Hayne | FAN | 60.68 | 7.83 | - | X | 0 | 0 | X | 0 | X |
| *Cratylia mollis* Mart. ex Benth. | FAN | 23.24 | 6.00 | - | X | 0 | 0 | X | 0 | X |
| *Dahlstedtia araripensis* (Benth.) M.J. Silva & A.M.G. Azevedo | FAN | 9.55 | 5.50 | - | 0 | 0 | 0 | X | 0 | 0 |
| *Dalbergia catingicola* Harms | FAN | 6.37 | 1.90 | - | 0 | 0 | 0 | X | 0 | 0 |
| Fabaceae 1 | FAN | 5.41 | 1.70 | - | 0 | 0 | 0 | X | 0 | 0 |
| Fabaceae 2 | FAN | 3.50 | 3.70 | - | 0 | 0 | 0 | X | 0 | 0 |
| *Mimosa hirsutissima* Mart. | CAM | - | - | 0.01 | 0 | 0 | 0 | X | 0 | - |
| *Mimosa ophthalmocentra* Benth. | CAM | - | - | 0.01 | 0 | 0 | 0 | X | 0 | - |
| *Mimosa* sp. | FAN | 3.58 | 3.25 | - | 0 | 0 | 0 | X | 0 | 0 |
| *Mimosa tenuiflora* (Willd.) Poir. | FAN | 26.74 | 3.33 | - | X | X | 0 | X | 0 | 0 |
| *Piptadenia stipulacea* (Benth.) Ducke | FAN | 25.06 | 5.83 | - | X | 0 | 0 | X | 0 | 0 |
| *Pityrocarpa moniliformis* (Benth.) Luckow & R. W. Jobson | FAN | 29.14 | 7.83 | - | 0 | 0 | 0 | X | 0 | 0 |
| *Poeppigia procera* C. Presl | FAN | 12.73 | 8.00 | - | 0 | 0 | 0 | 0 | 0 | 0 |
| *Cenostigma microphyllum* (Mart. ex G. Don) Gagnon & G.P. Lewis | FAN | 43.74 | 5.33 | - | X | 0 | 0 | X | 0 | 0 |
| *Cenostigma pyramidale* (Tul.) Gagnon & G.P. Lewis | FAN | 43.31 | 8.00 | - | X | 0 | 0 | 0 | 0 | X |
| *Senegalia piauhiensis* (Benth.) Seigler & Ebinger | FAN | 4.62 | 4.50 | - | X | 0 | 0 | X | 0 | 0 |
| *Senna martiana* (Benth.) H.S. Irwin & Barneby | FAN | 8.97 | 2.50 | - | X | 0 | 0 | X | 0 | 0 |
| *Senna rizzinii* H.S.Irwin & Barneby | CAM | - | - | 0.36 | 0 | 0 | 0 | X | 0 | - |
| *Senna splendida* (Vogel)H.S.Irwin & Barneby | CAM | - | - | 0.01 | 0 | 0 | 0 | X | 0 | - |
| *Stylosanthes seabrana* B. L. Maass & 't Mannetje | TER | - | - | 0.01 | 0 | 0 | 0 | X | X | - |
| *Trischidium molle* (Benth.) H.E. Ireland | FAN | 7.94 | 2.67 | - | X | 0 | 0 | X | 0 | X |
| *Zornia echinocarpa* (Meissner) Benth. | TER | - | - | 0.01 | 0 | 0 | 0 | X | X | - |
| **Lamiaceae** |  |  |  |  |  |  |  |  |  |  |
| *Hypenia salzmannii* (Benth.) Harley | TER | - | - | 6.28 | 0 | 0 | 0 | 0 | X | - |
| **Malpighiaceae** |  |  |  |  |  |  |  |  |  |  |
| *Barnebya harleyi* W.R. Anderson & B. Gates | FAN | 13.37 | 4.60 | - | X | 0 | 0 | 0 | 0 | 0 |
| *Byrsonima vacciniifolia* A. Juss. | FAN | 19.92 | 3.00 | - | X | 0 | 0 | 0 | 0 | X |
| *Galphimia brasiliensis* (L.) A.Juss. | TER | - | - | 0.01 | 0 | 0 | 0 | 0 | X | - |
| Malpighiaceae 1 | TER | - | - | 0.01 | 0 | 0 | 0 | 0 | X | - |
| Malpighiaceae 2 | TER | - | - | 0.34 | 0 | 0 | 0 | 0 | X | - |
| *Ptilochaeta* sp. | FAN | 8.22 | 6.07 | - | X | 0 | 0 | 0 | 0 | 0 |
| **Malvaceae** |  |  |  |  |  |  |  |  |  |  |
| *Herissantia tiubae* (K.Schum.) Brizicky | CAM | - | - | 4.97 | X | 0 | 0 | 0 | 0 | - |
| Malvaceae 1 | CAM | - | - | 0.01 | 0 | 0 | 0 | 0 | 0 | - |
| Malvaceae 2 | TER | - | - | 0.33 | 0 | 0 | 0 | 0 | 0 | - |
| Malvaceae 3 | TER | - | - | 0.33 | 0 | 0 | 0 | 0 | X | - |
| Malvaceae 4 | CAM | - | - | 0.67 | 0 | 0 | 0 | 0 | 0 | - |
| Malvaceae 5 | CAM | - | - | 0.66 | 0 | 0 | 0 | 0 | 0 | - |
| *Melochia* sp. | CAM | - | - | 7.02 | 0 | 0 | 0 | 0 | 0 | - |
| *Melochia tomentosa* L. | FAN | 5.49 | 2.50 | - | 0 | 0 | 0 | 0 | 0 | 0 |
| *Pavonia glazioviana* Gürke | FAN | 21.83 | 3.33 | - | X | 0 | 0 | 0 | 0 | 0 |
| *Pavonia luetzelburgii* Ulbr. | FAN | 8.91 | 1.40 | - | X | 0 | 0 | 0 | 0 | 0 |
| *Waltheria* sp. | FAN | 15.44 | 2.13 | - | 0 | 0 | 0 | 0 | 0 | 0 |
| *Waltheria* sp.2 | CAM | - | - | 1.33 | 0 | 0 | 0 | 0 | 0 | - |
| **Marantaceae** |  |  |  |  |  |  |  |  |  |  |
| *Maranta zingiberina* L.Andersson | TER | - | - | 0.01 | 0 | 0 | 0 | 0 | X | - |
| **Myrtaceae** |  |  |  |  |  |  |  |  |  |  |
| *Campomanesia eugenioides* (Cambess.) D.Legrand ex L.R. Landrum | FAN | 4.03 | 1.87 | - | 0 | 0 | 0 | 0 | 0 | X |
| *Psidium schenckianum* Kiaersk. | FAN | 5.73 | 2.00 | - | 0 | 0 | 0 | 0 | 0 | X |
| **Not identified** |  |  |  |  |  |  |  |  |  |  |
| Unidentified 1 | LIA | 11.93 | 5.00 | - | 0 | 0 | 0 | 0 | 0 | 0 |
| Unidentified 2 | FAN | 34.06 | 7.00 | - | 0 | 0 | 0 | 0 | 0 | 0 |
| Unidentified 3 | TER | - | - | 0.67 | 0 | 0 | 0 | 0 | X | - |
| Unidentified 4 | TER | - | - | 0.33 | 0 | 0 | 0 | 0 | X | - |
| Unidentified 5 | CAM | - | - | 0.66 | 0 | 0 | 0 | 0 | 0 | - |
| Unidentified 6 | TER | - | - | 0.01 | 0 | 0 | 0 | 0 | X | - |
| Unidentified 7 | TER | - | - | 0.01 | 0 | 0 | 0 | 0 | X | - |
| **Nyctaginaceae** |  |  |  |  |  |  |  |  |  |  |
| *Guapira tomentosa* (Casar.) Lundell | FAN | 29.68 | 6.50 | - | X | 0 | 0 | 0 | 0 | 0 |
| **Oxalidaceae** |  |  |  |  |  |  |  |  |  |  |
| *Oxalis divaricata* Mart. ex Zucc. | CAM | - | - | 0.01 | 0 | 0 | 0 | 0 | 0 | - |
| **Passifloraceae** |  |  |  |  |  |  |  |  |  |  |
| *Turnera diffusa* Willd. | FAN | 5.09 | 1.80 | - | 0 | 0 | 0 | 0 | 0 | 0 |
| **Phytolaccaceae** |  |  |  |  |  |  |  |  |  |  |
| *Microtea* *paniculata* Moq. | TER | - | - | 0.01 | 0 | 0 | 0 | 0 | X | - |
| **Plantaginaceae** |  |  |  |  |  |  |  |  |  |  |
| *Angelonia campestris* Nees & Mart. | CAM | - | - | 0.33 | X | 0 | 0 | 0 | 0 | - |
| *Tetraulacium veroniciforme* Turcz. | TER | - | - | 0.47 | 0 | 0 | 0 | 0 | X | - |
| **Poaceae** |  |  |  |  |  |  |  |  |  |  |
| *Axonopus* sp. | TER | - | - | 0.33 | 0 | 0 | 0 | 0 | X | - |
| *Digitaria* sp. | TER | - | - | 0.01 | 0 | 0 | 0 | 0 | X | - |
| *Digitaria* sp.2 | TER | - | - | 0.01 | 0 | 0 | 0 | 0 | X | - |
| *Eragrostis ciliaris* (L.) R.Br. | TER | - | - | 1.16 | 0 | 0 | 0 | 0 | X | - |
| *Panicum trichoides* Sw. | TER | - | - | 2.98 | 0 | 0 | 0 | 0 | X | - |
| *Setaria parviflora* (Poir.) M.Kerguelen | TER | - | - | 1.17 | 0 | 0 | 0 | 0 | X | - |
| *Setaria setosa* (Sw.) P.Beauv. | TER | - | - | 0.33 | 0 | 0 | 0 | 0 | X | - |
| *Urochloa* sp. | TER | - | - | 1.16 | 0 | 0 | 0 | 0 | X | - |
| **Polygalaceae** |  |  |  |  |  |  |  |  |  |  |
| *Asemeia ovata*(Poir.) J.F.B. Pastore & J.R. Abbott | TER | - | - | 0.67 | 0 | 0 | 0 | 0 | X | - |
| **Portulacaceae** |  |  |  |  |  |  |  |  |  |  |
| *Portulaca elatior* Mart. ex Rohrb. | TER | - | - | 0.01 | 0 | X | 0 | 0 | X | - |
| *Portulaca mucronata* Link | TER | - | - | 0.01 | 0 | 0 | 0 | 0 | 0 | - |
| *Portulaca oleracea* L. | HEM | - | - | 0.33 | 0 | 0 | 0 | 0 | 0 | - |
| **Rhamnaceae** |  |  |  |  |  |  |  |  |  |  |
| *Sarcomphalus joazeiro* (Mart.) Hauenschild | FAN | 5.57 | 1.40 | - | X | 0 | 0 | 0 | 0 | X |
| **Rubiaceae** |  |  |  |  |  |  |  |  |  |  |
| *Cordiera* sp.1 | FAN | 4.14 | 2.67 | - | X | 0 | 0 | 0 | 0 | X |
| *Cordiera* sp.2 | FAN | 9.30 | 1.50 | - | 0 | 0 | 0 | 0 | 0 | X |
| *Rubiaceae* 1 | TER | - | - | 0.33 | 0 | 0 | 0 | 0 | X | - |
| *Rubiaceae* 2 | TER | - | - | 0.33 | 0 | 0 | 0 | 0 | X | - |
| **Rutaceae** |  |  |  |  |  |  |  |  |  |  |
| *Balfourodendron molle* (Miq.) Pirani | FAN | 13.90 | 6.17 | - | 0 | 0 | 0 | 0 | 0 | 0 |
| **Sapindaceae** |  |  |  |  |  |  |  |  |  |  |
| *Cardiospermum corindum* L. | TER | - | - | 0.01 | 0 | 0 | 0 | 0 | X | - |
| Sapindaceae 1 | LIA | - | - | 0.01 | 0 | 0 | 0 | 0 | 0 | - |
| **Scrophulariaceae** |  |  |  |  |  |  |  |  |  |  |
| Scrophulariaceae | TER | - | - | 0.66 | 0 | 0 | 0 | 0 | X | - |
| **Selaginellaceae** |  |  |  |  |  |  |  |  |  |  |
| *Selaginella convoluta* (Arn.) Spring | HEM | - | - | 1.66 | 0 | 0 | 0 | 0 | 0 | - |
| **Simaroubaceae** |  |  |  |  |  |  |  |  |  |  |
| *Simaba ferruginea* A. St.-Hil. | FAN | 20.37 | 5.83 | - | 0 | 0 | 0 | 0 | 0 | 0 |
| **Solanaceae** |  |  |  |  |  |  |  |  |  |  |
| *Solanum agrarium* Sendtn. | CAM | - | - | 0.01 | 0 | 0 | 0 | 0 | 0 | - |
| *Solanum megalonyx* Sendtn. | FAN | 3.49 | 1.60 | - | 0 | 0 | 0 | 0 | 0 | 0 |
| *Solanum* sp. | TER | - | - | 0.01 | 0 | 0 | 0 | 0 | X | - |
| **Verbenaceae** |  |  |  |  |  |  |  |  |  |  |
| *Lantana camara* L. | FAN | 5.78 | 2.50 | - | 0 | 0 | 0 | 0 | 0 | 0 |
| *Lippia* sp. | FAN | 9.97 | 3.17 | - | X | 0 | 0 | 0 | 0 | 0 |
| *Lippia thymoides* Mart. & Schauer | FAN | 13.37 | 2.10 | - | X | 0 | 0 | 0 | 0 | 0 |
| Verbenaceae 1 | FAN | 10.19 | 1.40 | - | 0 | 0 | 0 | 0 | 0 | 0 |
| **Ximeniaceae** |  |  |  |  |  |  |  |  |  |  |
| *Ximenia americana* L. | FAN | 15.92 | 4.67 | - | X | 0 | 0 | 0 | 0 | X |

**Table S2:** Mean values of Functional Diversity (FRic - Functional Richness, FDiv - Functional Divergence, FEve - Functional Eveness and FDis - Functional Dispersion) considering all species and functional traits related to herbivory: resprouts, succulents with spines, urticancy/toxic and annual species. N=number of sample units. Low-disturbance regime (LOW) and the high-disturbance regime (HIGH).

|  | N | FRic | FDiv | FEve | FDis |
| --- | --- | --- | --- | --- | --- |
| NEAR | 4 | 17.03a | 0.86a | 0.14a | 1.31a |
| FAR | 4 | 13.74a | 0.91a | 0.08a | 1.69a |

**Table S3:** Net Relatedness Index (NRI) and Nearest Taxon Index (NTI) based on the unconstraint null model for the woody and non-woody component for the scale of 20-m x 50-m for woody and 20-m x 50-m for non-woody components. Significant p-value indicates that the phylogenetic structure differs from zero according to the t-test for one sample. N=number of plots of 20-m x 50-m for woody and 20-m x 50-m for non-woody. Low-disturbance regime (LOW) and the high-disturbance regime (HIGH)

|  |  | NRI |  |  | NTI |  |  |
| --- | --- | --- | --- | --- | --- | --- | --- |
|  | N | Mean | sd | p | Mean | sd | p |
| Woody component | | | | | | | |
| NEAR | 4 | -0.257 | 0.57 | 0.490 | 0.545 | 1.40 | 0.495 |
| FAR | 4 | 0.053 | 0.58 | 0.865 | -0.112 | 1.37 | 0.879 |
| Non-woody component | | | | | | | |
| NEAR | 4 | -0.013 | 0.68 | 0.971 | 1.242 | 1.37 | 0.168 |
| FAR | 4 | -0.513 | 0.34 | 0.055 | -0.650 | 1.27 | 0.380 |

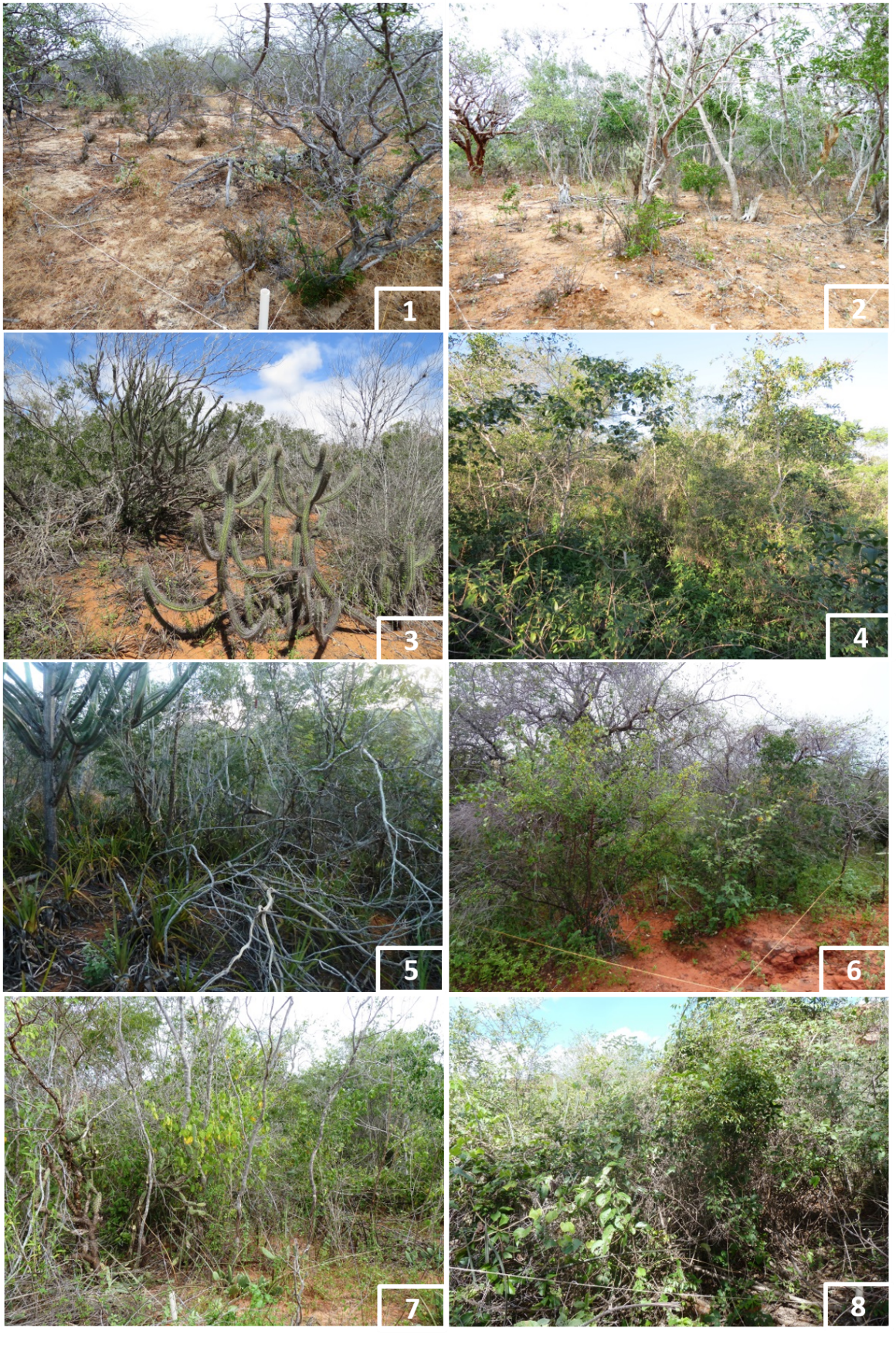

**Figure S1 -** Physiognomic aspect of the eight plots sampled in northern Bahia, Brazil. (1-4) Wood caatinga of the four plots near the road BR-235 that connects Juazeiro-BA to Carira-SE. (5-8) Wood caatinga of the four plots far from the road BR-235, Bahia. Photos: Mota, N. M. and Carrión, J. F.

| 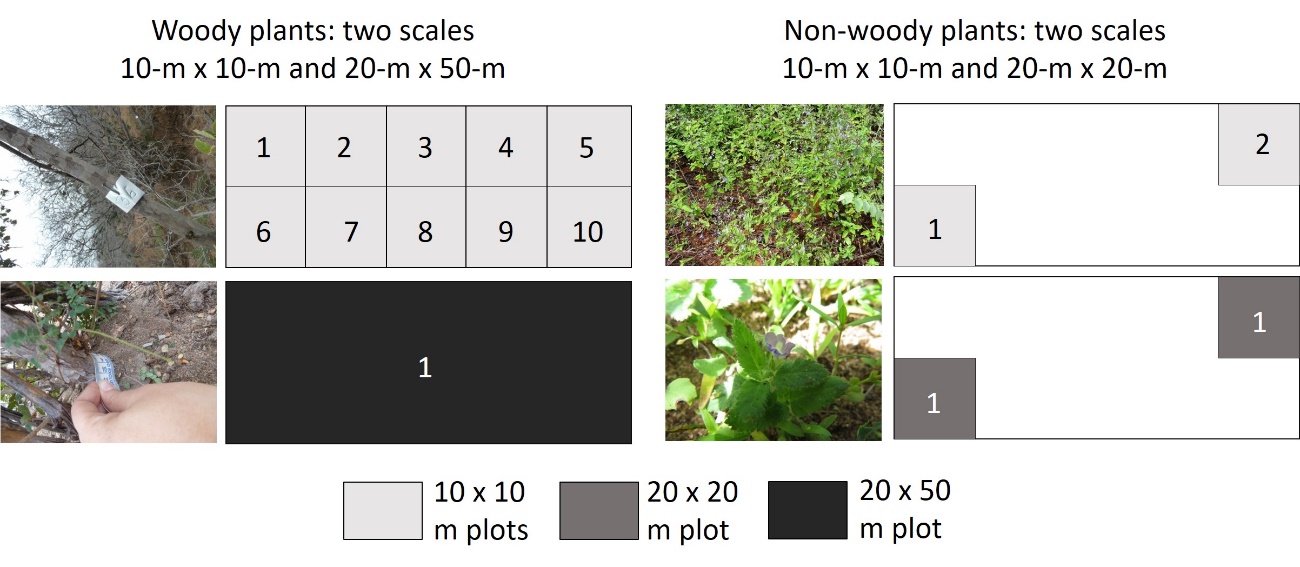 | 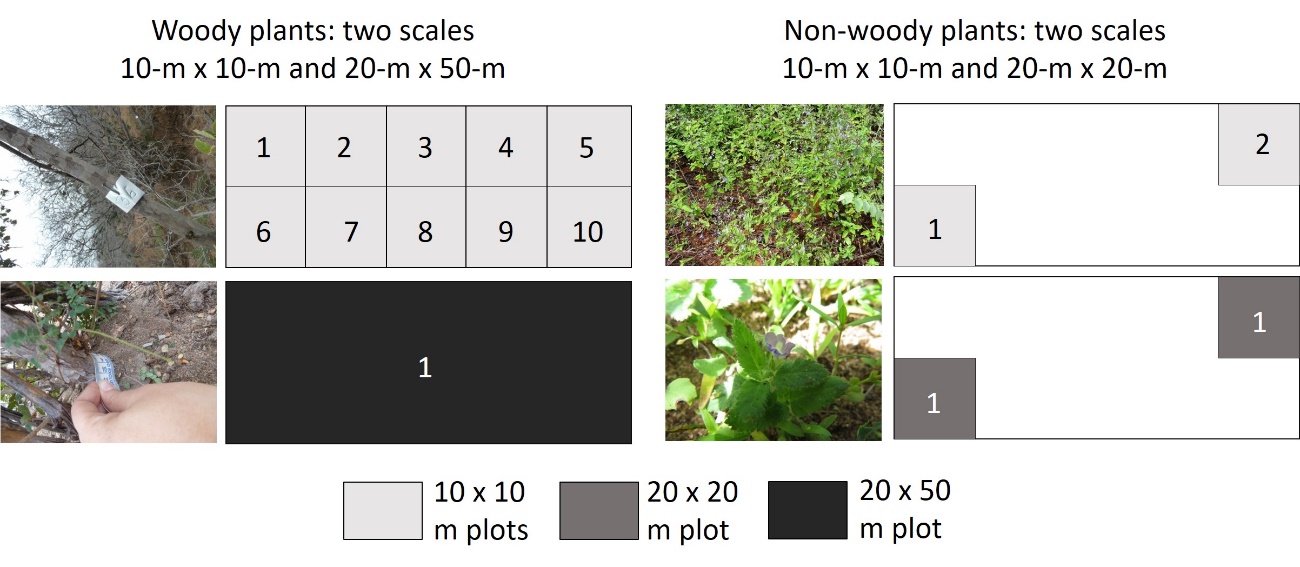 |
| --- | --- |

**Figure S2** – Scheme of plots and subplots of woody communities (left) at 10-m x10-m scale (top) and at 20-m x 50-m scale (bottom); scheme of plots and subplots of non-woody communities (right) at 10-m x 10-m scale and at 20-m x 50-m scale.

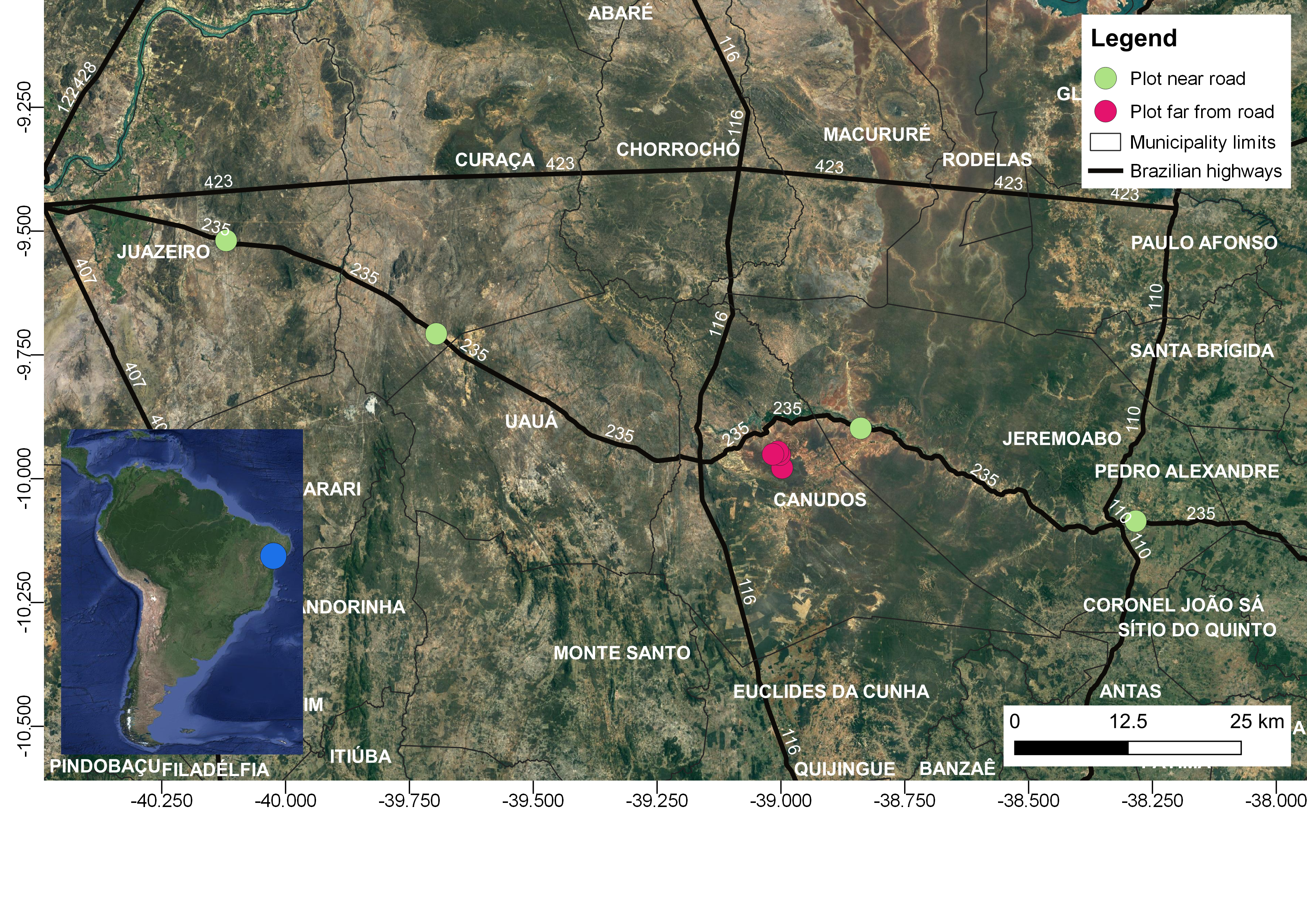

**Figure S3** - Road net and the samples. Thick lines are roads, thin lines are municipalities limits. Samples near road in three different municipalities (Juazeiro, Curaçá and Jeremoabo); samples further from roads in Canudos municipality with different situations of topography, and altitude. All samples are free of fences and domestic animals move freely.
